# Optimization of cerebrospinal fluid microbial metagenomic sequencing diagnostics

**DOI:** 10.1101/2020.06.25.170423

**Authors:** Josefin Olausson, Sofia Brunet, Diana Vracar, Yarong Tian, Sanna Abrahamsson, Sri Harsha Meghadri, Per Sikora, Maria Lind Karlberg, Hedvig Engström Jakobsson, Ka-Wei Tang

**Affiliations:** Department of Clinical Microbiology, Sahlgrenska University Hospital, Region Västra Götaland, Gothenburg, Sweden; Wallenberg Centre for Molecular and Translational Medicine, Department of Infectious Diseases, Institute of Biomedicine, University of Gothenburg, Gothenburg, Sweden; Clinical Genomics Gothenburg, Science for Life Laboratories, Gothenburg, Sweden; Department of Microbiology, Public Health Agency of Sweden, Solna, Sweden

**Keywords:** Metagenomics, Cerebrospinal fluid, Pathogen classification, PaRCA, Epstein-Barr virus

## Abstract

**Background:** Infection in the central nervous system is a severe condition associated with high morbidity and mortality. Despite ample testing, the majority of encephalitis and meningitis cases remain undiagnosed. Metagenomic sequencing of cerebrospinal fluid has emerged as an unbiased approach to identify rare microbes and novel pathogens. However, several major hurdles remains, including establishment of individual limits of detection, removal of false positives and implementation of universal controls.

**Results:** Twenty-one cerebrospinal fluid samples, in which a known pathogen had been positively identified by available clinical techniques, were subjected to metagenomic DNA sequencing using massive parallel sequencing. Fourteen samples contained minute levels of Epstein-Barr virus. Calculation of the detection threshold for each sample was made using total leukocyte content in the sample and environmental contaminants found in bioinformatic classifiers. Virus sequences were detected in all ten samples, in which more than one read was expected according to calculations. Conversely, no viral reads were detected in seven out of eight samples, in which less than one read was expected according to calculations. False positive pathogens of computational or environmental origin were readily identified, by using a commonly available cell control. For bacteria additional filters including a comparison between classifiers removed the remaining false positives and alleviated pathogen identification.

**Conclusions:** Here we show a generalizable method for detection and identification of pathogen species using metagenomic sequencing. The sensitivity for each sample can be calculated using the leukocyte count and environmental contamination. The choice of bioinformatic method mainly affected the efficiency of pathogen identification, but not the sensitivity of detection. Identification of pathogens require multiple filtering steps including read distribution, sequence diversity and complementary verification of pathogen reads.

## Background

Infections in the central nervous system (CNS) are severe and despite extensive microbiological diagnostic analysis a causative pathogen cannot be identified in many of the cases. A majority of CNS infections are caused by viruses, such as herpes simplex virus 1 (HSV1), varicella zoster virus (VZV or human herpesvirus 3) and enterovirus [1, 2]. Among CNS infections, *Streptococcus pneumoniae* and *Neisseria meningitidis* are the most common pathogens, while fungal or parasitic meningitis CNS infections are less common [3]. Epstein-Barr virus (EBV) has been implicated in recurrent meningitis and chronic encephalitis [4]. However, due to the high prevalence of EBV and its ability to remain latent in B-lymphocytes after primary infection and its role in tumorigenesis, assessing the clinical relevance of EBV DNA detected in cerebrospinal fluid (CSF) is difficult and presence of EBV is often considered to be an benign incidental finding [5, 6].

Current microbiological diagnostic methods include cultivation and nucleic acid detection of CSF, which are restricted to prior knowledge of the putative causing agent. Cultivation can detect a wide range of microorganisms, however, it is limited to viable and culturable pathogens. In contrast, nucleic acid detection is rapid and highly sensitive, but constrained to genetically conserved regions of known pathogens. Metagenomic sequencing using massive parallel sequencing, has the capability to discern multiple species and identify unknown species in samples. In metagenomics, the total nucleic acid present in the clinical sample is sequenced, thus provides an unbiased tool to diagnose infections and unknown species in samples [7-12].

Currently there is no standard for metagenomic sequencing in a clinical setting and the technique is still faced with some major challenges [13]. Contrary to PCR, the sensitivity in metagenomic sequencing is dependent on the fraction of pathogen sequences in the total sequencing library. Furthermore, laboratory contaminations detected in sequencing have been shown to differ greatly between laboratories and be dependent on the input biomass [14, 15]. Nucleic acid derived from the host and environmental contaminants must therefore be taken into account. Previous studies have calculated the sensitivity by using dilution of an exogenous pathogen into a known quantity of host background. However, this does not take into account the variability of clinical samples nor does it provide any guidance on how the sensitivity of each sample should be calculated.

Bioinformatic pathogen identification is a second major obstacle. Several publically available bioinformatic tools for classification are available, such as Centrifuge, Kraken and PathSeq [16-18]. Two conceptual different methods are frequently used, alignment of single reads (e.g. BLAST), or assemblies (k-mers), against pathogen databases. The list of pathogens generated by these applications are often long and requires exhaustive examinations in order to discern the true pathogen from bioinformatic misclassification and environmental contaminations. Criteria for identifying the causative pathogen include sequences disseminated throughout the microbial genome of the proposed pathogen, a threshold for number of pathogen reads in relation to total number of reads, and confirmation using several alignment algorithms have been suggested to increase the specificity [19, 20]. Each laboratory does however apply their own criteria.

We investigated the robustness of microbial metagenomics for clinical diagnostics of CNS-infections. To evaluate the diagnostic performance of the method, 21 CSF samples with variable levels of known pathogens were sequenced with the aim to identify factors important for calculating sensitivity. Also, four different taxonomic classifiers were assessed for their efficiency to identify pathogens as well as the number of false positive pathogens identified. Two commonly available cell lines were implemented as a positive and negative control to support the removal of environmental contaminants and bioinformatic misclassifications. Pathogen detection in DNA metagenomic sequencing in CSF is mainly limited by the leukocyte count which affects the sensitivity and bioinformatic missclassifications which affects the efficiency of pathogen identification.

## Results

We implemented a metagenomic DNA sequencing methodology to unbiasedly detect microbial species in CSF samples from patients with CNS symptoms in which a pathogen or EBV had been detected (Additional Table 1). Samples positively identified with pathogen-specific quantitative PCR (qPCR), 16S rRNA gene sequencing or bacterial/mycotic culture in CSF were included. Different pathogen types and variation of viral loads were chosen. CSF samples containing low levels of EBV were chosen to establish the sensitivity of the method. DNA from each sample was extracted and fragmented before library preparation and sequenced using massive parallel sequencing. Datasets were processed using five bioinformatic tools (Additional Figure 1).

**Table 1.**
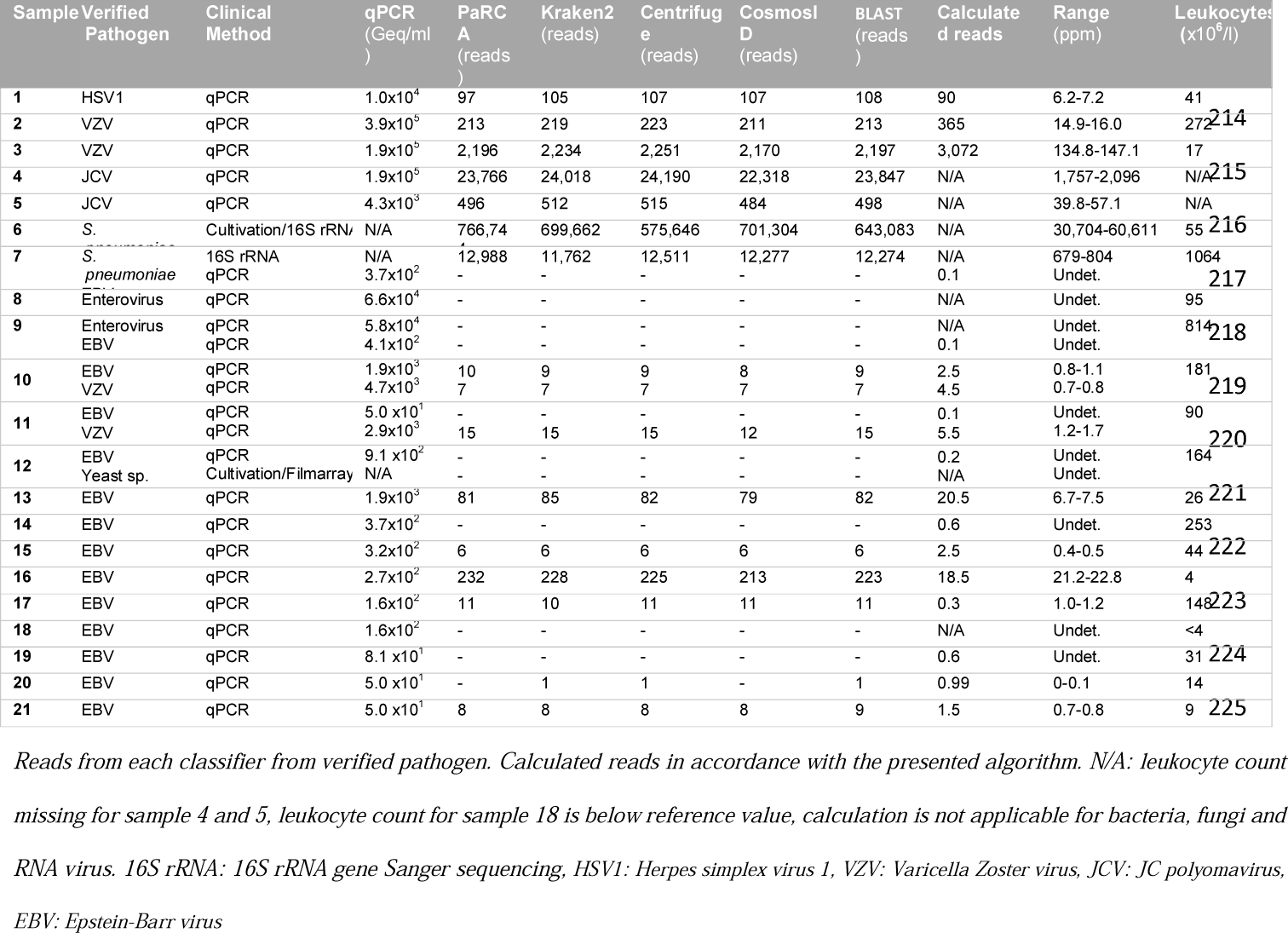
Metagenomic sequencing pipeline results.

**Figure 1.**
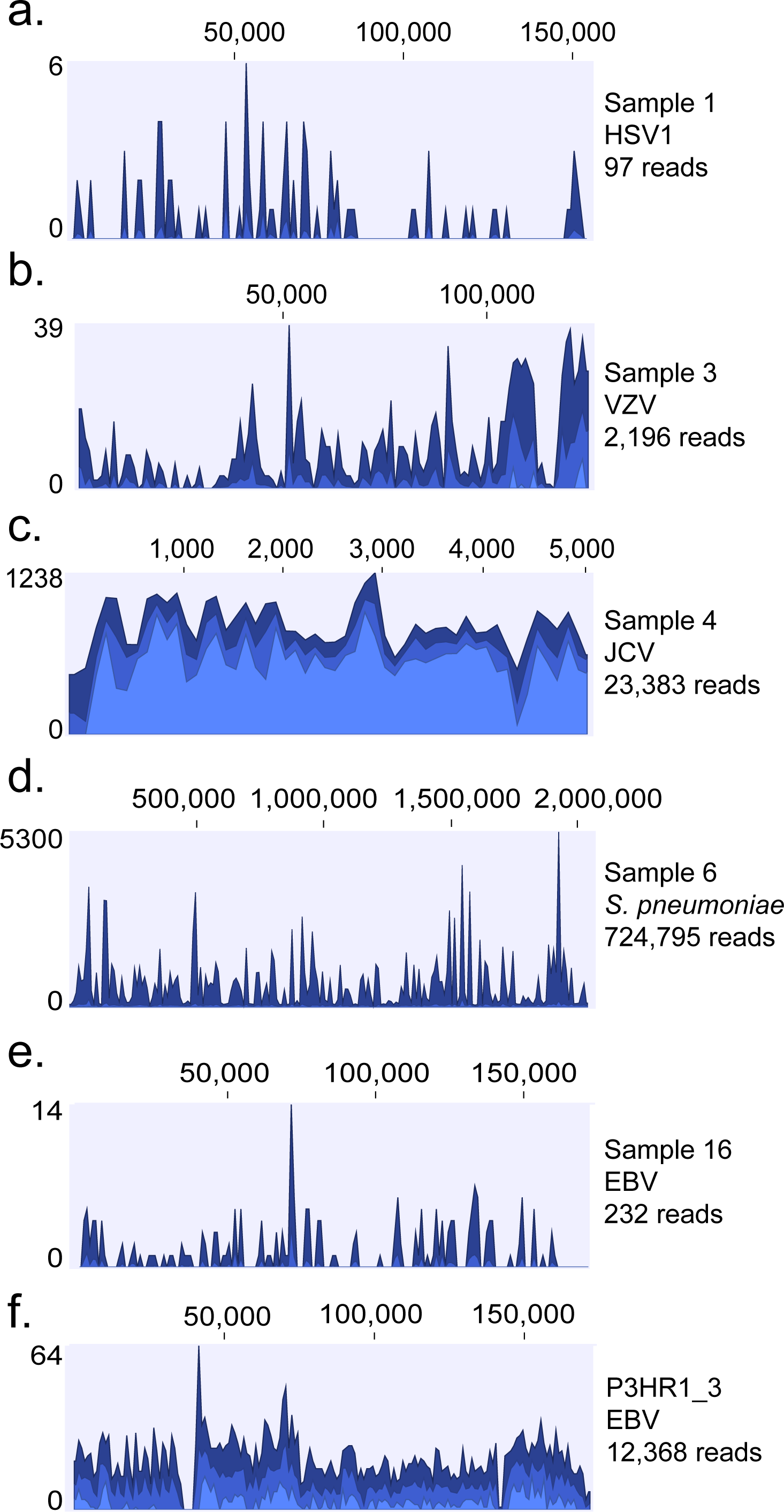
Pathogen genome alignment. Coverage density plot of sequencing reads from respective sample and control detected in PaRCA aligned to reference genomes of HSV1 (a), VZV (b), JCV (c), *S. pneumoniae* (d) and EBV (e-f). Number of reads (y-axis) at each nucleotide position of the genome (x-axis) depicted in blue. Dark blue represents peak, bright blue average and light blue minimum coverage for respective section of the genome.

### Bioinformatic classifiers

Four bioinformatic classifiers were included, Kraken2, Centrifuge, our in-house developed PaRCA (Pathogen detection for Research and Clinical Applications) and CosmosID. CosmosID was tested mainly for its ability to generate concise pathogen lists, but the format of the platform prevented a detailed analysis of the raw data and was therefore not included in all comparisons in the manuscript. The four bioinformatic classifiers diverged with regards to fraction of processed reads (from 85%-100%, Additional Table 2-3). However, the ability to identify the primary pathogen was similar comparing the classifiers.

### Sensitivity

Initially, three CSF samples (Sample 1-3) with high virus load of herpesvirus were analyzed. HSV1 and VZV were detected by all bioinformatic classifiers (Table 1). In sample 1, HSV1 was positively identified at 1×10^4^ genome equivalents per milliliter (Geq/ml) using qPCR. The sequencing library consisting of more than 15 million reads contained 6.2-7.2 HSV1 reads per million sequences analyzed (parts per million; ppm). The following two samples originated from patients with similar values of VZV DNA levels quantified by qPCR (1.9 and 3.9×10^5^ Geq/ml). Despite equivalent levels a ten-fold difference in detected VZV reads was observed between sample two (15-16 ppm) and sample three (135-147 ppm). Sample 2 contained 272×10^6^ white blood cells (WBC) per liter compared with sample 3 which contained 17×10^6^ WBC per liter (Table 1). We hypothesized that the difference in sensitivity was related to variations in leukocyte composition in the sample.

To further test the sensitivity, two CSF samples containing JC polyomavirus (JCV), a DNA virus with a relatively small genome, were processed. One sample contained high virus levels (1.9×10^5^ Geq/ml) and the other low virus levels (4.3×10^3^ Geq/ml) (Sample 4-5). JCV DNA was readily detected in both samples ranging from 1757-2096 ppm in sample 4 and 40-57 ppm in sample 5.

In order to verify that the methodology was applicable for bacterial agents, we sequenced CSF from two patients with pneumococcal meningitis, diagnosed by cultivation and/or 16S rRNA gene Sanger sequencing (Sample 6-7). DNA from *Streptococcus pneumoniae* (*S. pneumoniae*) was classified with a range between 30,704-60,661 ppm (Sample 6), and 679-804 ppm (Sample 7). In addition to the bacterial samples, we included two CSF samples from patients with RNA viral enterovirus CNS infection (Sample 8-9). As expected, no DNA reads were identified. Enterovirus was, however, found using metagenomic RNA sequencing (Additional figure 2)

**Figure 2.**
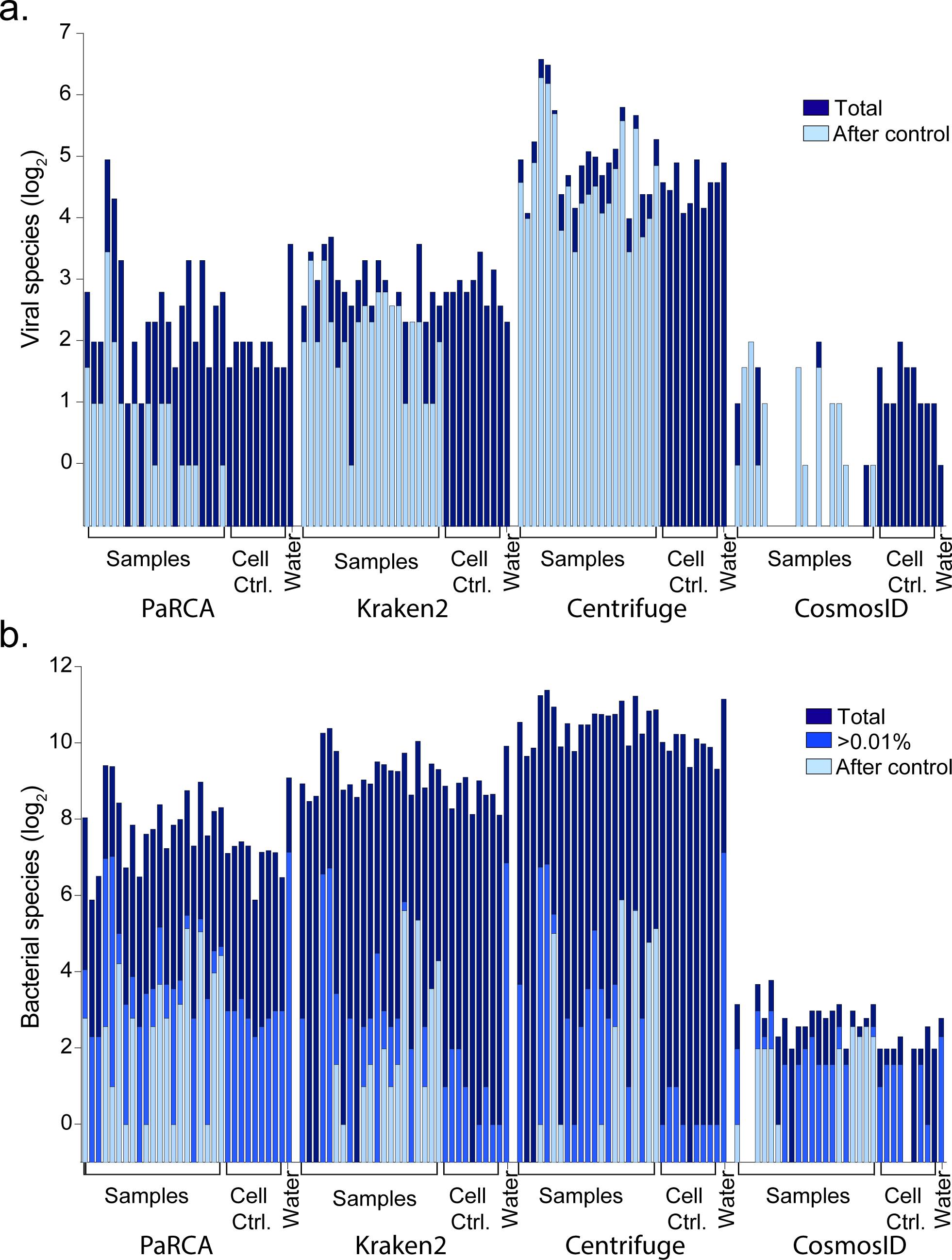
Detected pathogens in bioinformatic classifiers. Number of viral (a) and bacterial species (b) classified in each of the samples and controls using the different bioinformatic classifiers. Dark blue bars shows number of total number of species classified, bright blue bars shows amount of bacterial species over the fraction cutoff (≥0.01% of the dataset), light blue bars shows number of species not removed using controls.

Samples with co-infections, where EBV was detected along with a primary infectious agent (Enterovirus sample 9, VZV sample 10-11 and *Cryptococcus sp*. sample 12), were analyzed. Neither the EBV nor the enterovirus was detected in sample 9. VZV and EBV was detected in sample 10, and only VZV was detected in sample 11. Neither yeast nor EBV DNA was detected in sample 12. The results where expected when the following equation was applied for calculating the sensitivity for each agent.

The theoretically expected number of pathogen reads was calculated according to pathogen genome size (G_P_), the diploid human genome size of 6.5 billion basepairs (G_H_), pathogen copy according to PCR per milliliter (C_P_), whole blood cell count per milliliter (C_H_), and adjusted according to the volume (V), sequencing library size (L) and mappability in percent (M) to remove major contaminants.

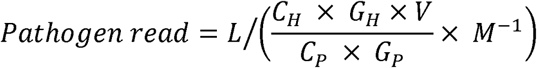

Thus, the detection limit of a single read of a pathogen with a 1 million basepair genome in CSF with normal WBC count (5×10^3^ per milliliter) using an input volume of 0.3 milliliter and >95% mappability require a sequencing library of approximately 10 million reads.

We included additional nine CSF samples with low levels of EBV DNA (50-2000 Geq/ml) (Sample 13-21). With the exception of sample 13 (patient diagnosed with CNS Hodgkin’s lymphoma type Post-Transplant Lymphoproliferative Disorder), and sample 16, where EBV was considered the cause of the symptoms, the EBV findings were clinically interpreted as benign incidental findings i.e. not the causative agent for the symptoms of infection. The EBV DNA detected in the majority of samples is likely to originate from latently infected B-lymphocytes recruited into the CSF. Despite the limitations for absolute quantification using qPCR and the stochasticity of distribution of low level pathogen particles, with one exception the calculated reads correlated with the detected reads in the sequencing data (Table 1). In ten samples, more than 1 viral reads was expected and pathogen sequences were found in all samples (Additional Figure 3). In seven samples where less than 1 read was expected to be found, EBV reads were only detected in one dataset (sample 17). Sixteen copies of EBV per milliliter was detected in sample 17 using qPCR and 11 reads were detected using metagenomic sequencing even though 0.3 reads were expected. The discrepancy between the calculation and and sequencing results is most likely due to the stochastic distribution of the few viral particles in the sample. In sample 20, 0.99 reads were expected to be detected in the dataset and a single EBV-read was identified in two of the four classifiers (Kraken2 and Centrifuge). This read was further confirmed using BLAST. The WBC count in sample 18 was below the reference interval of the leukocyte cell counter and was therefore omitted.

**Figure 3.**
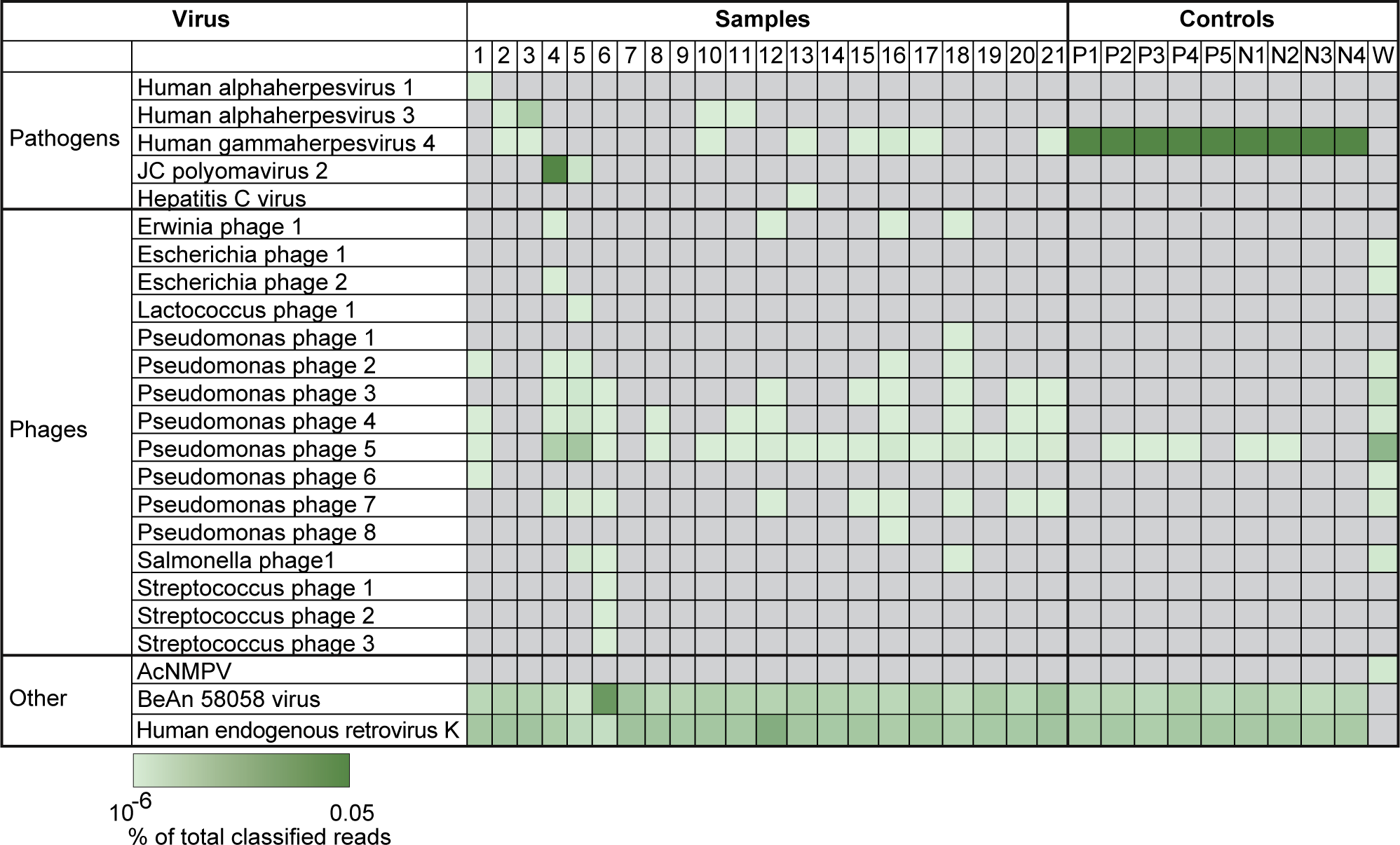
Viral species identified in datasets. Heatmap showing the ten most abundant viral species in each sample detected using PaRCA. AcMNPV: Autographa californica multiple nucleopolyhedrovirus. Controls: P; P3HR1, N; Namalwa, W; water.

All pathogen reads from PaRCA were mapped against the corresponding genome sequences using CLC genomics workbench (Figure 1a-e, Additional Figure 4). A dispersed distribution of the reads to the corresponding genomes was observed for all samples, except sample 10, where 5 of the 7 VZV reads (1 overlapping read) originate from a repetitive region within the genome and is therefore expected to be detected at a higher rate, and the last 2 reads map to a downstream gene (no overlap) (Additional Figure 4d). Each sequencing library was subjected to BLAST using the respective reference pathogen genome. The variation of the absolute number pathogen reads comparing the different classifiers detected was lower than 25% (Table 1).

**Figure 4.**
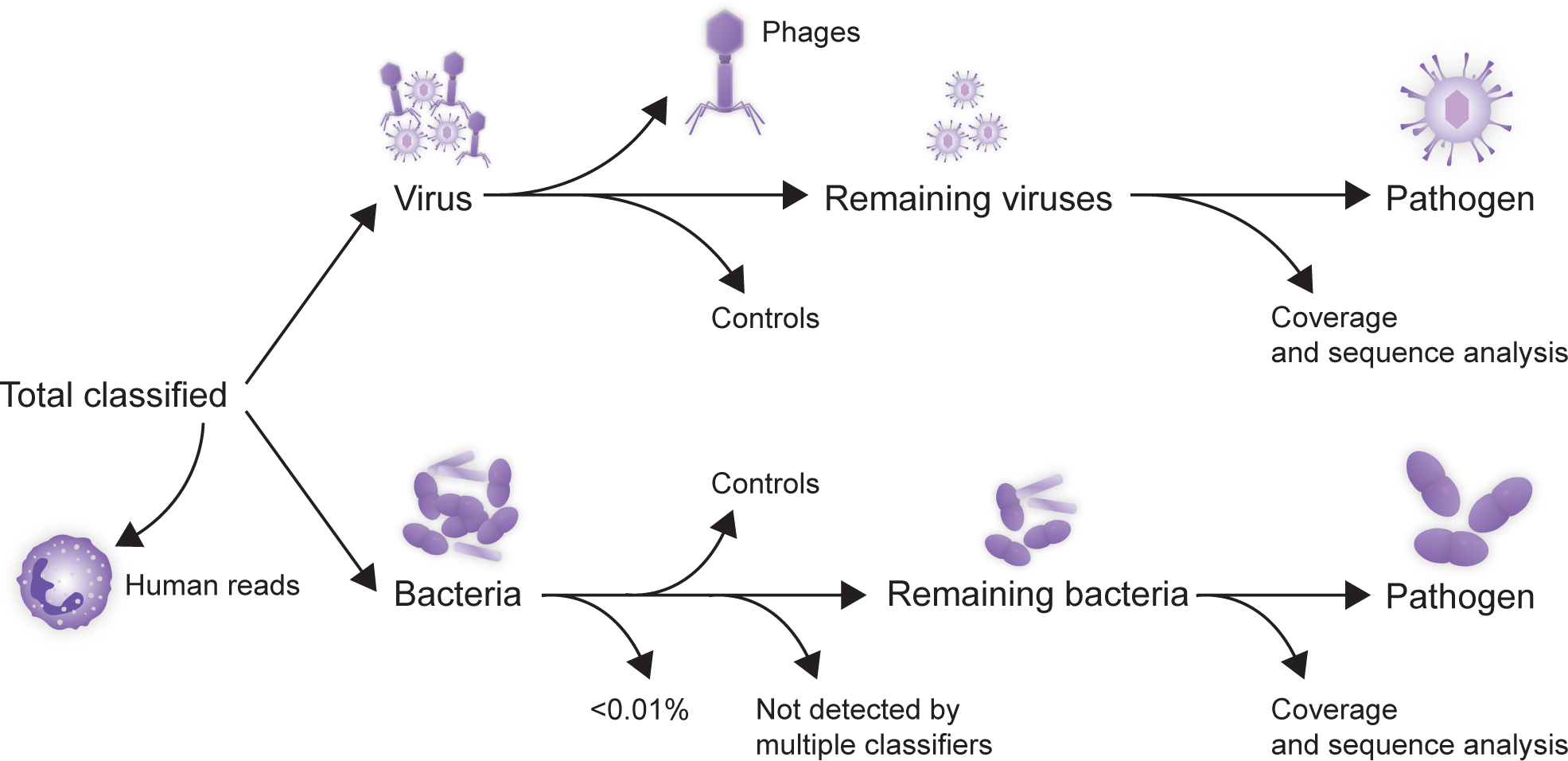
Discerning microbial pathogens from contaminations and misclassifications. Flowchart for identification of pathogens by removing false positive species. Virus contaminants can be removed by comparison of datasets with controls and manual examination of remaining viral reads. Phages can be disregarded as these virus do not infect human cells. Bacterial species require additional filters including a cutoff value and comparison between classifiers.

Qualitative and quantitative detection of a known pathogen can thus reproducibly be carried out using the different types of bioinformatic classifiers. Furthermore, an estimation of sensitivity for pathogens can be generated for each sample which can guide the clinician whether the sequencing depth is sufficient to find a certain type of pathogen (Additional Table 4). Notably however, each classifier produced diverse quantities of false positive hits.

### False positive pathogens

The diversity of viral species detected in metagenomic sequencing libraries were relatively low and recurrent. PaRCA, Kraken2, Centrifuge and CosmosID identified 2-31, 5-13, 17-96 and 0-4 viral species in each sample respectively (Figure 2a, Additional Table 5). Many of the most abundant viral species identified were found in multiple samples (Figure 3). Two samples (4 and 13) contained human virus which were not detected in multiple samples and not a previously confirmed pathogen (see below).

The non-pathogen/EBV viral reads were either of human origin, misclassified or contaminations. Human endogenous retrovirus K was identified in all samples, except for the water control, which was expected as the reads originates from the human genome (Figure 3 bottom, Additional Table 5). Another ubiquitously detected virus was the BeAN 58058 virus, which was detected in all samples, except for the water control. An additional BLAST examination identified these hits as human reads. Low levels of phage sequences known to infect bacteria from the *Enterobacterales* order were detected in a few samples and in the water control, most likely derived from bacteria purified enzymes used in the various steps of library preparation. A conspicuous pseudomonas phage contaminant in sample 4, 5 and the water control are likely derived from a bacterial contaminant at one of the sequencing sites. Streptococcus phage species were detected in sample 6, from a patient with *S. pneumoniae* meningitis. Importantly, the most prominent viral species identified in patient samples were also present in the cell controls at similar levels and displayed a similar sequence identity and could therefore be discarded as a pathogen.

Compared with the relatively few viral agents detected by the classifiers, bacterial species were abundant; 61-712 bacterial species were identified using PaRCA, 370-1408 in Kraken2, 845-2826 in Centrifuge and 0-14 in CosmosID (Figure 2b). Two samples originated from patients with a known *S. pneumoniae* meningitis (sample 6 and 7) and bacteria were detected at 69,088 ppm and 803 ppm resepectively (PaRCA). With the omission of the positive samples 6 and 7, trace levels (3.4-18.2 ppm) of *S. pneumoniae* was ubiquitously detected in all samples. A known environmental contamination of *Pseudomonas* was detected in the majority of the samples. In two samples (4 and 5) *Pseudomonas* constituted 389,480 ppm (39%) and 590,195 ppm (59%) of the entire sequencing library respectively, while the prevalence in other samples were lower 6.6-75,279 ppm (0.0007-7.5%). A large fraction of the detected bacteria are still left when using previously suggested fixed cut-off at 100 ppm (0.01%) (Figure 2) and unlike the virus species the contaminants/misclassifications cannot be entirely removed using the control samples. However, when further applying an additional filter of comparison of the detected bacterial species between the three classifiers (PaRCA, Kraken2 and Centrifuge) only the known pathogen (*S. pneumoniae*) or environmental contaminants (*Pseudomonas* and *Escherichia coli*) was left. Similarily no eukaryotic species were found in all three classifiers.

Considering the ubiquitous presence of viral misclassifications and contaminants in samples as well as controls, a viral pathogen is easily identifiable, but require additional analyses including read distribution and BLAST analysis, for verification in a clinical setting (discussed below). In contrast, the large number of bacterial species identified pose a bioinformatic challenge as the bacterial sequence can be derived from kit contaminants, lab environment or bioinformatic misclassifications which obscure the pathogen reads. As with the virus hits, removal of bacterial contaminants using cell controls can efficiently remove the majority of species, but additional filters are required (Figure 4).

### Controls

Two types of controls, water and cell control, were tested for their ability to mirror the bioinformatic missclassifications and contaminations observed in samples. In the water control the dataset consisted of 99.6% bacterial sequences and 0.06% viral sequences (Additional Table 5). The cell controls originating from EBV-transformed cancer cells had a composition more similar to the samples with 99.2-99.4% human sequences. The number of viral and bacterial strains detected in the water control was 12 and 568 respectively. In contrast the cell controls contain sequences ranging from 3-4 viral and 61-177 bacterial strains.

The viral strains in the water control were mainly of phage origin. In contrast the viral strains detected in the cell controls were similar to the CSF samples, mainly Human endogenous retrovirus K and BeAN 58058 virus. Both cell lines originate from EBV-transformed cancer cells and harbours EBV DNA. The ppm-values of each cell line between sequencing runs was reproducible and no significant difference was found between the classifiers (Additional Figure 5, Additional Table 6).

In the water control, 98% of the sequencing library consisted of reads from *Pseudomonas* and the second most abundant bacterial strain found was *Escherichia coli* (0.1%), which is to be expected as most enzymes are produced in this bacterial system. In contrast, none of the bacterial strains in the cell controls constituted more than 0.1% of the sequencing library.

Thus, the water control efficiently amplified the environmental and kit contaminants, but in contrast to the cell control did not find human misclassifications. Also, since the water control consist entirely of contaminants, the absolute or proportionate content did not allow for a direct comparison with the patient samples. The cell control allowed for direct quantitive and qualitative subtraction of the majority of contaminants and putative pathogens were identified.

### Unexpected virus findings

In sample 2 and 3 we identified 29-34 EBV reads in both samples in all classifiers (Additional Table 5). The reads were dispersed throughout the genome and displayed minor sequence variability with the reference genome in accordance with previous EBV findings (Additional Figure 6a-b). Due to the limited sample volume we were unable to verify and quantify this finding using qPCR.

In sample 4 we identified three viruses which were unexpected, mastadenovirus, papillomavirus and torque teno virus (Additional Figure 6c-e). PaRCA identified 32 reads matching human mastadenovirus C (HAC), Kraken2 32 reads, Centrifuge 30 reads and CosmosID did not report any HAC sequences. The majority of reads, 25 out of 32 where 198 bp long, 5 reads where shorter and 2 were longer. BLAST-analysis showed that all reads shared the same 3’-end. Four reads had mismatches in comparison with reference sequence. Considering the size and distribution of the reads our findings are most likely a laboratory amplicon contamination. Human papillomavirus (HPV) reads were detected in PaRCA (12 reads), Kraken2 (2 reads), but not by Centrifuge and CosmosID. Ten of the 12 reads were 105 bp long and the remaining two, 104 bp and 106 bp respectively. All reads aligned to the 3’-end of the virus genome in the L1 gene. Examination of BLAST results showed a high similarity with HPV98 with a one or two base-pair mismatch. As above, considering the size and distribution of the reads our findings were most likely a laboratory amplicon contamination. CosmosID has an inbuilt function to filter out hits that are considered to be amplicons, therefore the software did not report these reads. Different strains of Anellovirus/Torque teno virus (TTV) were detected in the classifiers. PaRCA identified 75 reads, Kraken2 25 reads, Centrifuge 55 reads, while CosmosID did not detect any TTV reads. Five distinct consensus reads/contigs were formed from the 75 reads identified in PaRCA. Thirty-one reads formed a consensus reads of 196bp. BLAST analysis of this read displayed a 97% identity with TTV14, but only for 91bp of the fragment. The remaining parts of the contig did not show any alignment with any viral species. The origin of this read is therefore unknown. BLAST analysis of the remaining 4 reads/contigs showed alignment (>95% query cover and identity) to an Anellovirus isolate previously identified in metagenomics. The alignment showed an unusual coverage of the 5’-end of the genome and all the reads were aligned to the first half of the genome. The reason for this unusual coverage is unknown, but considering that TTV is widely detected in metagenomic sequencing and the multiple reads aligning to a clinical isolate it is probable that these four contigs/reads originate from the patient sample.

In sample 13, we detected 10 reads corresponding to hepatitis C virus (HCV) in PaRCA. Kraken2, Centrifuge and CosmosID detected 5, 6 and 6 reads respectively. The 10 reads were concentrated to the 5’-end of the genome, but spread within the initial half of the genome (Additional Figure 6f). An analysis of the BLAST results showed alignment with HCV genotype 1. Synonymous mutations were found in multiple reads as well as gaps. Two reads had a fusion between sequences from different regions of the HCV genome. The sequence diversity indicates that the virus is from a patient, but the frameshift and fusion reads indicates that they are of an artificial origin. Also, the patient had undergone HCV serology analysis which was negative. Finally, considering that HCV is a RNA virus this finding is most likely a laboratory amplicon contamination.

## Discussion

In this study we subjected 21 CSF samples from patients with suspected or confirmed CNS infection to metagenomic DNA sequencing. Pathogen detection accuracy and efficiency was evaluated using five bioinformatic tools. Using 12 samples with minute levels of EBV we concluded that the sensitivity of detection was mainly affected by leukocyte content in the samples and to lesser degree environmental contamination. Bioinformatic classifiers were essentially equally efficient in terms of sensitivity, but produced vastly different number of false positive hits, which inhibited efficient clinical pathogen identification. The removal of these false positive hits originating from contaminants and bioinformatics classifications were alleviated by using a EBV-containing cell control which served as a positive as well as a negative control. A number of criteria have been suggested for how to identify a causative agent in clinical samples e.g. by calculating the fraction of pathogen reads and/or an absolute number of reads. However, using these methods the majority of samples used in this study would be considered negative and/or contain a large number of agents which would be considered falsely positive dependent on the choice of classifier. The lower detection limit could be generalized and compared between studies/laboratories if the leukocyte count was provided. In a similar manner, a general quantification of viral content using ppm is an efficient reference point for comparison between studies [21, 22]. Furthermore, it is evident that local contaminants greatly impact the sequencing library constitution. Therefore, it is necessary that findings in negative controls from each study is presented in its entirety. Nine CSF-samples were identified at the clinic to only contain EBV, and we did not identify any additional pathogen, confirming the results from the clinic. Importantly, using our algorithm a lower detection limit could be determined for pathogens. An alternative to metagenomic sequencing is removal of the dominating host background using various methods including centrifugation and nuclease treatment [23, 24]. However, this will deplete the majority of nucleic acid and only minute amounts of nucleic acid will be left, which complicates the library preparation. Sensitivity would also be reduced, especially for intracellular virus, and bacteria which might precipitate if centrifugation is used. Likewise the specificity would be impaired by the overwhelming number of environmental contaminants as seen in our water control.

Our bioinformatic classifier PaRCA, which uses a combination of single reads alignment and assemblies was able to detect more reads from HAC, HPV, TTV and HCV, but failed to detect the single EBV read in sample 20. Bioinformatic classifiers for clinical practice should not only quantify the pathogen reads, but also provide information of read distribution, sequence diversity and subtraction of environmental contaminants and bioinformatic misclassifications, facilitating pathogen detection as shown in this study. Novel pathogens will also require classifiers to detect diverse sequences, as well as enable investigation of sequences which might not classify completely to a genus. Our finding of a novel TTV strain shows that there is a large difference between bioinformatics classifiers ability to identify divergent sequences.

In this study we have used archived material, which impair a proper RNA analysis due to degradation. Future studies using fresh CSF-samples where RNA integrity and quantity is measured may provide similar guidelines for RNA pathogen detection. We only included two verified bacterial CSF-samples in this study, one which was detected by culturing and 16S rRNA gene sequencing, and the second one detected by 16S rRNA gene sequencing. A limit of using metagenomic sequencing of CSF from bacterial meningitis patients is the high levels of leukocytes, but this may be compensated by the higher amount of bacterial nucleic acid compared with viral genomes. Here, we applied a fraction cutoff for bacterial findings (>0.01%) in order to decrease the amount of false positive bacterial species findings. This cutoff value should not be considered fixed and future studies with larger bacterial cohort would provide additional guidelines for bacterial species identification.

## Conclusions

We suggest that prior to clinical metagenomic DNA sequencing, an estimation of sequencing depth is made by adjusting it to the leukocyte content in the sample. Also, a pathogen-containing cell control sequenced at the same depth should be included in the same sequencing run in order to generate the same type of reproducible background. Bioinformatic processing should include a comparison between the pathogens detected in the cell control and the sample as well as between multiple classifiers. Further candidate pathogens reads should be confirmed by using BLAST and mapped against a reference genome to identify read distribution and sequence diversity. A comprehensive evaluation including a theoretical estimation on sensitivity of the metagenomics test as well as other clinical microbiological assays e.g. serology and PCR should assist the clinician in interpreting the final results.

## Methods

### Sample collection

Included in this retrospective study were cerebrospinal fluid samples from patients with CNS symptoms of infection, in which the Department of Clinical Microbiology at Sahlgrenska University Hospital or the The Public Health Agency of Sweden previously had verified the infectious agent during 2015-2017. The sample cohort was chosen to include a variety of microorganisms (DNA/RNA virus, bacteria or fungi) with varying concentration of the pathogens as determined by confirmatory testing using qPCR, cultures, 16S rRNA gene Sanger sequencing or FilmArray (Additional Methods). The samples were stored at −20°C after clinical testing. The cell lines P3HR1 (HTB-62, American Type Culture Collection, ATCC, USA) and Namalwa (CRL-1432, American Type Culture Collection, ATCC, USA), were used as combined negative controls as well as positive controls, due to its inherent EBV genome. The controls were processed in parallel with the patient samples during all the laboratory steps.

### Sample processing

For samples processed at the Department of Clinical Microbiology at Sahlgrenska University Hospital, total nucleic acid was extracted from 400 µl of cerebrospinal fluid using the MagNA Pure Compact Nucleic Acid Isolation Kit I (Roche Diagnostics, Indianapolis, IN, USA) on the MagNA Pure compact automated extractor. For samples processed at The Public Health Agency of Sweden, total nucleic acid was extracted from 200 µl of cerebrospinal fluid sample using the MagDEA® Dx SV (Precision System Science Co Ltd, Matsudo-city, Chiba, Japan) on the magLEAD® 12gC automated extractor (Precision System Science Co Ltd). DNA concentrations were determined using the Qubit Fluorometer (Thermo Fisher Scientific, Waltham, MA, USA) using the dsDNA HS Assay Kit (Thermo Fisher).

### Library preparation and sequencing

DNA libraries were prepared according to the modified protocol for metagenomic samples, developed at the Public Health Agency of Sweden, using the Ion Xpress Plus Fragment Library Kit (Thermo Fisher) on the AB Library Builder System (Thermo Fisher). Samples were fragmented to 200 bp, followed by ligation of Ion P1 Adapter as well as Ion Xpress Barcode adapters. The protocol was adjusted to suit low-input samples (<50 ng DNA) by using a reduced volume of P1 adapter and barcodes (0.5 µl). The libraries were amplified, selecting the number of amplification cycles according to the sample input concentration, varying between 14 to 20 cycles. Amplified libraries were size selected choosing an optimal size range for each individual sample to ensure removal of small-sized PCR concatemers, varying between 100 to 320 bp (including adapters). Size selection was performed using the Pippin Prep platform (Sage Science, Beverly, MA, USA) with 2% Dye free Agarose Gel Cassette. Following visualization and an estimation of the concentration using the High Sensitivity D1000 DNA Kit on the Agilent 2200 TapeStation system (Agilent Technologies, CA, USA), the samples were pooled according to concentration. Subsequently, libraries were purified using Agencourt AMPure XP (Beckman Coulter, Brea, CA, USA). Finally, libraries were quantified using qPCR with the Ion Library TaqMan Quantitation Kit (Thermo Fisher) and the size estimated using High Sensitivity D1000 DNA Kit on Agilent 2200 TapeStation system (Agilent Technologies). For template preparation, libraries were pooled to a final concentration of 50 pM, if obtainable. For libraries with lower concentration than 50 pM, libraries were pooled to the available concentration. Thereafter, the Ion Chef Platform was used to ligate the libraries onto spheres using the Ion 540 Kit-Chef (Thermo Fisher). Following clonal amplification, libraries were loaded onto Ion 540 Chip and sequencing was performed on the S5 System (XL, Prime; Thermo Fisher) according to the manufacturer’s protocol for 200 bp read length.

### Bioinformatic analysis

#### Quality Control

BAM-files were converted into fastq files using the Torrent Suite Software provided for Ion S5 system. Reads were processed with FASTX toolkit [25] to fasta files. Fastqc was used to identify low-quality reads. Sequences were then subjected to the individual pipelines described below.

### Pathogen detection for Research and Clinical Applications (PaRCA)

Databases were created using built-in tools in Kraken2 and Kaiju. Briefly, databases, corresponding to bacteria, viruses and eukaryotes were created at DNA, RNA and protein level resulting in nine total k-mer databases. The viral databases were comprised of all viral data in GenBank, the bacterial database consisted of the full Progenomes data [26] and eukaryotic databases were composed of the GenBank data for vertebrates, parasites and fungi.

After download, the Progenomes database was continuously updated using scripts to reflect changes within the NCBI taxonomy. Reads were initially trimmed at both directions using BBDuk (BBMap 37.50) using an entropy mask of 0.9, trim quality of 16 and a minimum length of 40. Reads were corrected using Fiona (0.2.9) with id=3 for substitution errors.

Reads were classified using Kraken2 and Kaiju by using individual databases. Kraken2 results were filtered using the kraken-filter with a threshold of 0.15 for eukaryotes and 0.05 for viruses and bacteria (a higher threshold indicates higher stringency). Thresholds for Kaiju: score and minimum matches were set to 85.20 for eukaryotes, 80.18 for bacteria and 75.15 for viruses.

After initial classification and filtering, Kraken2 results were individually compared and reads with hits in multiple databases were evaluated based on k-mer score with the highest scoring match being retained for further downstream analysis. Kaiju scores were internally compared and the hit with the longest protein alignment was preserved. Reads with both Kraken2 and Kaiju hits were then compared and the lowest common ancestor of the two results was selected using mergeOutputs with “-c lowest” from the Kaiju package. Reads where the lowest common ancestor was a species designation were directly counted and saved while reads with a higher lowest common ancestor were further processed in the pipeline. Reads only classified by a single k-mer classifier were labelled as “singletons” and further processed.

Reads were ordered by taxonomic ID, which then were regressed through the taxonomic tree until either a genus-level or kingdom-level was reached. Reads without genus-level information or reads with a classification above genus level were stored separately for further analysis. After ordering into genus, all taxonomic IDs corresponding to a member of the genus were automatically downloaded from NCBI and corresponding accession identifiers were parsed from the NCBI accession dump file. Accession identifiers were then used to create a slice of the BLAST nt-database for that specific genus. Reads classified as belonging to the order “primates” was not processed further and received the taxonomic ID 9606 (Homo Sapiens).

Reads were analyzed in BLAST within the genus using a threshold of an e-value of 10^-3^ and the ten best hits were then retained. The ten results per read were parsed and the bit-score per taxon in the hits were aggregated. The taxon with the highest aggregate bit score was then selected as the putative taxon ID for the read. After taxon identification, results were merged and regressed in order to identify the species level classification of the putative taxon. If the kingdom level was reached before a species identification was found, the original taxon identifier was used in its place. Finally, any reads that were not successfully classified within a genus in the BLAST database creation step were collected and subjected to BLAST against the full NT-database with an e-value of >10^-5^ and a minimum query coverage of 20% as threshold, again the ten best hits were preserved. The results from both BLAST analyses were aggregated based on bit score and the resulting taxon ID regressed to species level if possible.

Classified reads were collected and presented using a krona-graph and tables in an html format. Tables were reorderable on name, taxonomic id and read count. Tables were also filterable, including wildcard functionality. FASTQ-files containing reads classified to an individual species and aggregates corresponding to kingdoms and unclassified reads were directly downloadable.

### Kraken2

We used Kraken2 with a dustmasker included in the package.

### Centrifuge

We subjected our samples to Centrifuge with the inbuilt quality control and repeatmasker based on dustmasking from NCBI tools. Briefly, the dustmasker converts the low-quality regions into N’s so the aligner skips aligning these sequences [16]. In order to obtain reads from all pathogens included in this study, the total of both leaf and genus levels were incorporated from the Centrifuge reports, thus leading to higher amounts of total classified reads, however, since not all species were converted into the ETE3 toolkit, and some stops at genus level, this does not affect final results of classified pathogens.

### CosmosID

Unassembled sequencing reads were directly analyzed using the commercially available genomic platform CosmosID to achieve identification of microbes at species level [27]. Each uploaded sample was searched and cleared from host sequences by the platform prior to analysis. CosmosID automatically filters out phages and amplicon-originated sequences.

### BLAST

BLAST analysis was performed with reference genomes for the pathogens. The cutoff was set to ≥95% sequence identity and an e-value of ≤10^-3^. Following standard steps for pre-processing reads, a BLAST search was performed with reads set as subjects and reference genomes set as queries. Reference genomes used were NC_001806 (HSV1), NC_001348 (VZV), NC_00196 (JCV), NC_003098 (*S. pneumoniae*), NC_007605 (EBV), NC_001405 (Human Mastadenovirus C; MAVC), FM_955837.2 (Human Papillomavirus 98; HPV98), MH_649255.1 (Anellovirus), and NC_004102.1 (HCV).

### Calculations and statistical analysis

CLC genomics workbench (Ver. 11, Qiagen) was used to perform and plot coverage analysis. Classified sequences from Kraken2 and Centrifuge were visualized using Pavian [28]. Ratio between sample ppm and control ppm were calculated, where an ratio ≤ 10 were considered a contamination.

GrapPad Prism Ver. 7.0c was utilized to perform statistical analysis. Kruskal-Wallis test with Dunn’s multiple comparison tests was applied to compare reproducibility through pipelines. A *p*-value ≤ 0.05 were considered significant.

## Declarations

### Ethics approval and consent to participate

The study design and methods were approved by the Regional ethical review board in Gothenburg (191-18).

### Availability of data and materials

Will be available on the European Genome-phenome Archive upon publication.

### Author contributions

This study was designed by HEJ, MLK, SB and KWT. Cells were cultured by YT. Samples were selected by KWT, SB and DV. Metagenomic sequencing was performed by MLK, SB, and JO. PS, SA and SHM executed bioinformatic analysis. Calculations was done by JO, SA and KWT. DV and KWT provided clinical expertise. Manuscript was written by JO, SB, HEJ and KWT. Figures and tables was prepared by JO, YT and KWT. All authors read and approved the final manuscript.

## Acknowledgements

We thank the Bioinformatics Core Facility at the Sahlgrenska Academy for bioinformatics analyses.

## Additional information

**Additional table 1**. Clinical data (docx)

**Additional table 2**. Dataset species classification (docx)

**Additional table 3**. Pathogen detection by bioinformatic classifier (docx)

**Additional table 4**. Patient report with estimation of sensitivity for pathogens (xlsx)

**Additional table 5**. Species identified in bioinformatic classifiers (xlsx)

**Additional table 6**. Cell control reproducibility (docx)

**Additional method**. Description of clinical methodology used for comparison (docx)

**Additional figure 1**. Overview of the sample and bioinformatic processing (pdf)

DNA from cerebrospinal fluid specimens was extracted and followed by library construction and sequencing. Datasets generated by the Ion S5 were processed by four different bioinformatics classifiers to profile the microbiome. BLAST was used for verification.

**Additional figure 2**. Enterovirus samples (pdf)

Results of viral species detected in RNA sequencing datasets of sample 8 and sample 9 in PaRCA.

**Additional figure 3**. Correlation between detected and calculated reads (pdf)

The reads detected by PaRCA correlated to the calculated reads using the algorithm, Spearman Correlation coefficient, n=10

**Additional figure 4**. Coverage density plot of microbial species in CSF samples (pdf)

Reads from samples not shown in main figure mapped to reference genomes of (a) VZV (NC_001348), (b) JCV (NC_00196), (c) *S. pneumoniae* (NC_003098), (d, f) VZV (NC_001348), and (e, g-j) EBV (NC_007605) using CLC Genomics Workbench. Number of reads (y-axis) at each nucleotide position of the genome (x-axis) depicted in blue. Dark blue represents peak, bright blue average and light blue minimum coverage for respective section of the genome.

**Additional figure 5**. Cell control coverage density plot and reproducibility (pdf)

Coverage analysis of EBV reads detected in cell controls Namalwa (a) and P3HR1 (b) mapped to EBV reference genome (NC_007605) using CLC Genomics Workbench. Number of reads (y-axis) at each nucleotide position of the genome (x-axis) depicted in blue. Dark blue represents peak, bright blue average and light blue minimum coverage for respective section of the genome. EBV reads shown as parts per million reads (ppm) in each of the cell line controls for each of the bioinformatic classifier (c), n=4 (Namalwa) or n=5 (P3HR1); Kruskal-Wallis test with Dunn’s multiple comparisons show no significant difference between the pipelines.

**Additional figure 6**. Coverage analysis for unexpected findings (pdf)

Reads from samples with ambigous findings mapped to reference genomes of EBV NC_007605 (a-b), Human Mastadenovirus C (MAVC) NC_001405 (c), Human Papillomavirus 98 (HPV98) FM_955837.2 (d), Anellovirus MH_649255.1 (e), and HCV NC_004102.1 (f), using CLC Genomics Workbench.

