## Additional Table 1 for "Optimization of cerebrospinal fluid microbial metagenomic sequencing diagnostics"

**Additional Table 1.** Clinical data.

| Sample | Age/Sex | Diagnosis | Verified pathogen | Confirmatory testing | CSF cell count (×10^6^/l) | | | | | Clinical Data | |
| --- | --- | --- | --- | --- | --- | --- | --- | --- | --- | --- | --- |
|  |  |  |  | (Geq/ml) | Ery | Lym | Mon | NeNeu | Pol | |  |
| 1 | 60/F | Encephalitis | HSV1 | qPCR (1.0x10^4^) | 1 |  | 40 |  | 1 | | Fever, dizziness, fainting, confusion, convulsions, pathological Romberg’s sign |
| 2 | 32/M | Meningitis | VZV | qPCR (3.9x10^5^) | 2 | 254 | 18 | 0 |  | | Fever, dizziness, headache, photosensitivity, nausea, vomiting, common cold symptoms, blisters dorsal and ventral left flank, diplopia left eye, pathological Romberg’s sign, neck stiffness. |
| 3 | 24/M | Meningitis | VZV | qPCR (1.9x10^5^) | 92 | 17 | 0 | 0 |  | | Secretory otitis media ongoing healing, common cold symptoms, headache, vomiting |
| 4 | N/A | N/A | JC polyomavirus | qPCR (1.9x10^5^) | N/A | | | | | | N/A |
| 5 | N/A | N/A | JC Polyomavirus | qPCR (4.3x10^3^) | N/A | | | | | | N/A |
| 6 | 65/F | Meningitis  Acute Mastoiditis, Pneumoniae | *S. pneumoniae* | Cultivation and 16S rRNA gene Sanger Seq | 5405 |  | 10 |  | 45 | | Fever, unconsciousness, sore ears, petechiae-lower extremities, positive Babinski’s sign, neck stiffness |
| 7 | 55/M | Meningitis, Otitis media | *S. pneumoniae*  EBV | 16S rRNA gene Sanger seq qPCR (3.7x10^2^) | 112 | 76 | 265 | 723 |  | | Fever, dizziness, headache, common cold symptoms, obtundation, confusion, perforated otitis media, sunset glance, enlarged pupil, weakness on left side, neck stiffness |
| 8 | 37/M | Viral meningitis | Enterovirus | qPCR (6.6x10^4^) | <5 | 65 | 26 | 4 |  | | Headache, photosensitivity, common cold symptoms, fainting, pathological Romberg's sign, pathological finger-nose test |
| 9 | 11/M | Meningoencephalitis | Enterovirus EBV | qPCR (5.8x10^4^) qPCR (4.1x10^2^) | 13 |  | 484 |  | 330 | | Fever, dizziness, headache, photosensitivity, nausea, memory loss, difficulty feeling touch – lower leg |
| 10 | 83/F | Encephalitis | EBV VZV | qPCR (1.9x10^3^) qPCR (4.7x10^3^) | 106 | 160 | 19 | 2 |  | | Fever, constitutional symptoms, varicella zoster, right hemiparesis, neck stiffness |
| 11 | 86/F | Shingles (Orthostatism) | EBV VZV | qPCR (5.0 x10^1^) qPCR (2.9x10^3^) | 27 | 67 | 23 | <3 |  | | Dizziness, nausea, initial fluctuating confusion, varicella zoster right leg, pathological finger-nose test |
| 12 | 57/F | Meningitis | EBV Yeast sp. | qPCR (9.1 x10^2^) Cultivation and Filmarray | 13340 |  | 161 |  | 3 | | Dizziness, headache, blurred vision, falling tendency, dragging walk, weakness left leg, pathological Romberg’s sign |
| 13 | 52/F | CNS Hodgkin's lymphoma | EBV | qPCR (1.9x10^3^) | 5 | 26 | <3 | <3 |  | | Dizziness, nausea, vomiting, constitutional symptoms, somnolence, left hemiparesis, dysarthria |
| 14 | 5/M | Neuroborreliosis | EBV | qPCR (3.7x10^2^) | 403 | 178 | 49 | 26 |  | | Fever, headache, nausea, vomiting, sore neck, previous foul bite without rash |
| 15 | 67/M | Viral encephalitis | EBV | qPCR (3.2x10^2^) | <5 | 32 | 12 | <3 |  | | Confusion, dysphasia with paraphasia and neologisms, pathological finger-nose test |
| 16 | 14/M | Encephalitis, Staphylococcal sepsis, Viral pneumoniae, T-ALL | EBV | qPCR (2.7x10^2^) | <5 | 4 | <3 | <3 |  | | Fever, constitutional symptoms, confusion, slurred speech |
| 17 | 30/M | Postoperative neurosurgical infection | EBV | qPCR (1.6x10^2^) | 220 |  | 300 |  | 148 | | Fever, dizziness, headache, nausea |
| 18 | 74/M | No infection | EBV | qPCR (1.6x10^2^) | 3 | <3 | <3 | 0 |  | | Headache, nausea, vomiting, constitutional symptoms, neck stiffness |
| 19 | 21/F | Encephalitis | EBV | qPCR (8.1 x10^1^) | <5 | 31 | <3 | <3 |  | | Fever, dizziness, headache, common cold symptoms, nausea, vomiting, confusion, convulsions (mycoplasma serum IgM positive and nasopharyngeal swab PCR positive) |
| 20 | 88/F | Encephalitis | EBV | qPCR (5.1 x10^1^) | <5 | 14 | <3 | <3 |  | | Fever, dizziness, nausea, vomiting, constitutional symptoms, depersonalization, wide-range walk, tendency falling backwards, dysarthria, neck stiffness |
| 21 | 72/K | Neuromyelitis optica | EBV | qPCR (5.0 x10^1^) | 91 |  | 8 |  | 1 | | Dermatitis, leg paresis, dissociated feeling, impairment, weakness |

*Geq: Genome equivalents, Ery: erythrocytes, Lym: lymphocytes, Mon: monocytes, Neu: neutrophils, Pol: polycytes. HSV1: Herpes simplex virus 1, VZV: Varicella Zoster virus, EBV: Epstein-Barr virus, T-ALL: T-cell acute lymphoblastic leukemia*
