## Additional Table 2 for "Optimization of cerebrospinal fluid microbial metagenomic sequencing diagnostics"

**Additional Table 2.** Datasets species classification

| Sample |  | | PaRCA reads | | | | Kraken2 reads | | | | | Centrifuge reads* | | | | CosmosID hits^#^ | | | | |
| --- | --- | --- | --- | --- | --- | --- | --- | --- | --- | --- | --- | --- | --- | --- | --- | --- | --- | --- | --- | --- |
|  | Total reads | Classified | | Human | Virus | Bacteria | Classified | Human | Virus | Bacteria | Classified | | Human | Virus | Bacteria | | Classified | Human | Virus | Bacteria |
| 1 | 16,155,106 | **15,240,521** | | 15,101,081 | 4,372 | 195,129 | **15,859,247** | 15,706,386 | 117 | 150,227 | **17,250,776** | | 17,064,379 | 149 | 174,726 | | **16,155,106** | N/A | 226 | 59,618 |
| 2 | 14,278,116 | **13,280,177** | | 13,232,349 | 4,431 | 24,125 | **13,820,286** | 13,817,757 | 259 | 605 | **14,862,675** | | 14,850,476 | 269 | 1,262 | | **14,275,166** | N/A | 356 | 1,413 |
| 3 | 16,041,251 | **14,924,255** | | 14,859,234 | 6,755 | 36,199 | **15,522,840** | 15,512,386 | 2,273 | 1,526 | **16,699,082** | | 16,673,106 | 2,321 | 2,131 | | **16,041,251** | N/A | 2,393 | 1,910 |
| 4 | 12,703,224 | **11,338,709** | | 6,797,953 | 29,530 | 4,475,933 | **11,728,272** | 7,143,611 | 24,242 | 4,577,006 | **13,348,386** | | 7,928,803 | 25,421 | 5,284,937 | | **12,703,224** | N/A | 25,986 | 1,754,615 |
| 5 | 12,165,130 | **10,525,017** | | 4,231,556 | 5,576 | 6,265,762 | **10,923,115** | 4,498,521 | 878 | 6,422,262 | **12,451,198** | | 4,948,147 | 1,656 | 7,294,357 | | **12,165,130** | N/A | 2,887 | 2,549,867 |
| 6 | 14,957,135 | **12,650,219** | | 11,396,742 | 6,437 | 1,153,776 | **14,486,711** | 13,517,631 | 11 | 963,584 | **18,748,448** | | 17,194,162 | 100 | 1,526,405 | | **14,957,135** | N/A | 1,275 | 104,057 |
| 7 | 17,365,318 | **16,152,332** | | 16,069,006 | 6,081 | 50,834 | **16,923,847** | 16,904,670 | 7 | 16,840 | **18,419,777** | | 18,389,723 | 22 | 25,307 | | **17,365,318** | N/A | 171 | 4,012 |
| 8 | 13,417,152 | **12,532,220** | | 12,383,644 | 3,657 | 121,529 | **13,134,090** | 13,046,745 | 8 | 84,987 | **14,371,346** | | 14,261,140 | 35 | 98,431 | | **13,417,152** | N/A | 149 | 33,851 |
| 9 | 11,006,034 | **10,128,505** | | 10,086,217 | 2,791 | 25,089 | **10,634,909** | 10,634,909 | 9 | 4,147 | **11,408,918** | | 11,400,094 | 20 | 5,319 | | **11,006,034** | N/A | 94 | 2,555 |
| 10 | 9,600,578 | **8,934,196** | | 8,859,290 | 2,840 | 59,202 | **9,322,094** | 9,277,062 | 24 | 242,141 | **10,019,619** | | 9,962,846 | 43 | 50,731 | | **9,600,578** | N/A | 113 | 17,385 |
| 11 | 9,569,552 | **8,900,918** | | 8,792,316 | 2,620 | 93,511 | **9,283,282** | 9,204,476 | 20 | 77,768 | **9,984,472** | | 9,885,888 | 51 | 89,799 | | **9,535,729** | N/A | 123 | 30,691 |
| 12 | 12,244,759 | **11,413,586** | | 10,932,187 | 4,997 | 409,339 | **11,919,736** | 11,566,102 | 26 | 342,404 | **13,796,034** | | 13,377,407 | 82 | 379,079 | | **12,244,759** | N/A | 202 | 133,832 |
| 13 | 11,791,941 | **10,831,390** | | 10,701,867 | 3,548 | 110,867 | **11,387,096** | 11,291,000 | 98 | 94,739 | **12,260,830** | | 12,143,605 | 115 | 107,741 | | **11,791,941** | N/A | 239 | 36,766 |
| 14 | 15,668,502 | **14,591,084** | | 14,460,476 | 4,308 | 107,011 | **15,213,606** | 15,128,599 | 10 | 83,995 | **16,264,830** | | 16,259,511 | 35 | 97,425 | | **15,668,502** | N/A | 149 | 34,593 |
| 15 | 13,852,556 | **12,977,595** | | 12,790,541 | 3,836 | 164,700 | **13,499,122** | 13,356,211 | 19 | 141,659 | **14,487,022** | | 14,303,887 | 49 | 163,618 | | **13,809,049** | N/A | 172 | 56,224 |
| 16 | 11,061,515 | **10,165,144** | | 9,347,773 | 3,159 | 798,825 | **10,678,731** | 9,880,623 | 241 | 769,897 | **11,604,819** | | 10,670,040 | 351 | 918,003 | | **11,061,515** | N/A | 511 | 315,290 |
| 17 | 9,993,372 | **9,264,766** | | 9,217,416 | 2,976 | 30,329 | **9,690,977** | 9,673,204 | 15 | 16,641 | **10,417,701** | | 10,294,763 | 27 | 19,439 | | **9,993,372** | N/A | 101 | 7,717 |
| 18 | 19,727,930 | **18,336,989** | | 17,070,118 | 4,779 | 1,234,359 | **19,159,304** | 17,942,646 | 27 | 1,215,263 | **20,761,857** | | 19,342,338 | 127 | 1,391,846 | | **19,727,930** | N/A | 480 | 499,185 |
| 19 | 9,611,223 | **8,942,217** | | 8,867,132 | 2,858 | 60,693 | **9,347,387** | 9,296,863 | 5 | 49,591 | **10,034,970** | | 9,972,248 | 22 | 57,726 | | **9,611,223** | N/A | 111 | 19,746 |
| 20 | 11,375,823 | **10,526,555** | | 10,242,680 | 2,955 | 264,731 | **11,061,201** | 10,818,124 | 10 | 242,141 | **11,927,331** | | 11,640,347 | 24 | 270,426 | | **11,375,823** | N/A | 147 | 95,885 |
| 21 | 10,687,870 | **10,016,450** | | 9,709,562 | 3,836 | 289,961 | **10,398,272** | 10,108,840 | 17 | 288,096 | **11,207,425** | | 10,869,174 | 68 | 327,877 | | **10,687,870** | N/A | 211 | 119,483 |

** The total of both leaf and genus levels were merged from the Centrifuge reports, leading to higher amounts of total classified reads.*

*^#^ Human reads were not provided by CosmosID, hits instead of reads*
