## Additional Table 3 for "Optimization of cerebrospinal fluid microbial metagenomic sequencing diagnostics"

**Additional Table 3.** Pathogen detection by bioinformatic classifier.

| Sample |  |  |  | PaRCA | | Kraken2 | | Centrifuge* | | CosmosID | | BLAST |
| --- | --- | --- | --- | --- | --- | --- | --- | --- | --- | --- | --- | --- |
|  | Verified  Pathogen | Method  (Geq/ml) | Total reads Ion Torrent | Reads/total class. reads | ppm | Reads/total class. reads | ppm | Reads/total class. reads | ppm | Reads/total  class. reads | ppm | Unique Reads |
| 1 | HSV1 | qPCR (1.0x10^4^) | 16,155,106 | 97/ 15,240,521 | 6.4 | 105/ 15,859,247 | 6.6 | 107/ 17,250,776 | 6.2 | 107/ 16,155,106 | 6.6 | 108 |
| 2 | VZV | qPCR (3.9x10^5^) | 14,278,116 | 213/ 13,280,177 | 16.0 | 219/ 13,820,286 | 15.9 | 223/ 14,862,675 | 15.0 | 211/ 14,275,166 | 14.9 | 213 |
| 3 | VZV | qPCR (1.9x10^5^) | 16,041,251 | 2,196/ 14,924,255 | 147.1 | 2,234/ 15,522,840 | 143.9 | 2,251/ 16,699,082 | 134.8 | 2,170/ 16,041,251 | 135.3 | 2197 |
| 4 | JCV | qPCR (1.9x10^5^) | 12,703,224 | 23,766/ 11,338,709 | 2,096.0 | 24,018/ 11,728,272 | 2,047.9 | 24,190/ 13,348,386 | 1,812.2 | 22,318/12,703,224 | 1,756.9 | 2, 847 |
| 5 | JCV | qPCR (4.3x10^3^) | 12,165,130 | 496/ 10,525,017 | 47.1 | 512/ 10,923,115 | 46.9 | 515/ 12,451,198 | 41.4 | 484/ 12,165,130 | 39.8 | 498 |
| 6 | *SP* | Cultivation & 16S rRNA  gene Seq | 14,957,135 | 766,744/ 12,650,219 | 60,611.1 | 699,662/ 14,486,711 | 48,296.8 | 575,646/ 18,748,448 | 30,703.7 | 701,304/ 14,957,135 | 46,887.6 | 643,083 |
| 7 | *SP*  EBV | 16S rRNA gene seq  qPCR (3.7x10^2^) | 17,365 318 | 12,988/ 16,152,332  0/ 16,152,332 | 804.1  - | 11,762/ 16,923,847  0/ 16,923,847 | 695.0  - | 12,511/ 18,419,777  0/ 18,419,777 | 679.2  - | 12,277/ 17,365,318  0/ 17,365,318 | 707.0  - | 12,274 0 |
| 8 | EV | qPCR (6.6x10^4^) | 13,417,152 | 0/ 12,532,220 | - | 0/ 13,134,090 | - | 0/ 14,371,346 | - | 0/ 13,417,152 | - | 0 |
| 9 | EV  EBV | qPCR (5.8x10^4^)  qPCR (4.1x10^2^) | 11,006,034 | 0/ 10,128,505  0/ 10,128,505 | -  - | 0/ 10,634,909  0/ 10,634,909 | -  - | 0/ 11,408,918  0/ 11,408,918 | -  - | 0/ 11,006,034  0/ 11,006,034 | -  - | 0 0 |
| 10 | EBV  VZV | qPCR (1.9x10^3^)  qPCR (4.7x10^3^) | 9,600,578 | 10/ 8,934,196  7/ 8,934,196 | 1.1  0.8 | 9/ 9,322,094  7/ 9,322,094 | 1.0  0.8 | 9/ 10,019,619  7/ 10,019,619 | 0.9  0.7 | 8/ 9,600,578  7/ 9,600,578 | 0.8  0.7 | 9 7 |
| 11 | EBV  VZV | qPCR (5.0 x10^1^)  qPCR (2.9x10^3^) | 9,569,552 | 0/ 8,900,918  15/ 8,900,918 | -  1.7 | 0/ 9,283,282  15/ 9,283,282 | -  1.6 | 0/ , 984,472  15/ 9,984,472 | -  1.5 | 0/ 9,535,729  12/ 9,535,729 | -  1.2 | 0 15 |
| 12 | EBV  Yeast | qPCR (9.1 x10^2^)  Cultivation & Filmarray | 12,244,759 | 0/11,413,586  0/11,413,586 | -  - | 0/ 11,919,736  0/ 11,919,736 | -  - | 0/ 13,796,034  0/ 13,796,034 | -  - | 0/ 12,244,759  0/ 12,244,759 | -  - | 0 0 |
| 13 | EBV | qPCR (1.9x10^3^) | 11,791,941 | 81/ 10,831,390 | 7.5 | 85/ 11,387,096 | 7.5 | 82/ 12,260,830 | 6.7 | 79/ 11,791,941 | 6.7 | 82 |
| 14 | EBV | qPCR (3.7x10^2^) | 15,668,502 | 0/ 14,591,084 | - | 0/ 15,213,606 | - | 0/ 16,264,830 | - | 0/ 15,668,502 | - | 0 |
| 15 | EBV | qPCR (3.2x10^2^) | 13,852,556 | 6/ 12,977,595 | 0.5 | 6/ 13,499,122 | 0.4 | 6/ 14,487,022 | 0.4 | 6/ 13,809,049 | 0.5 | 6 |
| 16 | EBV | qPCR (2.7x10^2^) | 11,061,515 | 232/ 10,165,144 | 22.8 | 228/ 10,678,731 | 21.4 | 225/ 11,604,819 | 19.4 | 213/ 11,061,515 | 21.2 | 223 |
| 17 | EBV | qPCR (1.6x10^2^) | 9,993,372 | 11/ 9,264,766 | 1.2 | 10/ 9,690,977 | 1.0 | 11/ 10,417,701 | 1.1 | 11/ 9,993,372 | 1.1 | 11 |
| 18 | EBV | qPCR (1.6x10^2^) | 19,727,930 | 0/ 18,336,989 | - | 0/19,159,304 | - | 0/ 20,761,857 | - | 0/ 19,727,930 | - | 0 |
| 19 | EBV | qPCR (8.1 x10^1^) | 9,611,223 | 0/ 8,942,217 | - | 0/ 9,347,387 | - | 0/ 10,034,970 | - | 0/ 9,611,223 | - | 0 |
| 20 | EBV | qPCR (5.1 x10^1^) | 11,375,823 | 0/ 10,526,555 | - | 1/ 11,061,201 | 0.1 | 1/ 11,927,331 | 0.1 | 0/ 11,375,823 | - | 1 |
| 21 | EBV | qPCR (5.0 x10^1^) | 10,687,870 | 8/ 10,016,450 | 0.8 | 8/ 10,398,272 | 0.8 | 8/ 11,207,425 | 0.7 | 8/ 10,687,870 | 0.8 | 9 |

*Reads/total class. reads: Pathogen reads/total reads classified by bioinformatical pipeline. ppm: parts per million, abundance of pathogen per millions of total classified reads.*

*JCV: JC polyomavirus, SP: Streptococcus pneumonia, EV: Enterovirus*

**The total of both leaf and genus levels were incorporated in the Centrifuge reports, leading to higher amounts of total classified reads, however this did not affect the final amount of pathogen reads.*
