## Additional Table 6 for "Optimization of cerebrospinal fluid microbial metagenomic sequencing diagnostics"

**Additional Table 6.** Cell control reproducibility.

|  |  | **PaRCA** | | | **Kraken2** | | | **Centrifuge** | | | **CosmosID** | | |
| --- | --- | --- | --- | --- | --- | --- | --- | --- | --- | --- | --- | --- | --- |
| Control | ID# | Classified reads | EBV Reads | ppm (mean±SD) | Classified reads | EBV Reads | ppm (mean±SD) | Classified reads | EBV Reads | ppm (mean±SD) | Classified reads | EBV Reads | ppm (mean±SD) |
| **Namalwa**  **Cell line** | **#1** | 14,441,248 | 16,584 | **1006±225** | 15,085,293 | 16,894 | **955±199** | 16,136,669 | 16,952 | **920±210** | 15,561,865 | 16,040 | **913±201** |
|  | **#2** | 11,757,402 | 14,115 |  | 12,208,229 | 13,227 |  | 13,008,822 | 14,341 |  | 12,505,333 | 13,672 |  |
|  | **#3** | 14,385,091 | 10,095 |  | 15,021,622 | 10,232 |  | 16,081,341 | 10,212 |  | 15,325,051 | 9,828 |  |
|  | **#4** | 7,490,706 | 7,280 |  | 7,788,053 | 7,289 |  | 8,291,564 | 7,387 |  | 7,970,628 | 7,063 |  |
| **P3HR1**  **Cell line** | **#1** | 12,641,680 | 14,798 | **1335±207** | 13,260,465 | 15,133 | **1210±183** | 14,189,956 | 15,155 | **1215±192** | 13,693,487 | 14,265 | **1191±176** |
|  | **#2** | 9,483,576 | 12,753 |  | 9,847,965 | 12,924 |  | 10,520,818 | 12,930 |  | 10,060,502 | 12,307 |  |
|  | **#3** | 11,421,831 | 12,410 |  | 12,029,195 | 12,665 |  | 12,962,700 | 12,677 |  | 12,388,273 | 12,010 |  |
|  | **#4** | 9,191,839 | 13,973 |  | 9,634,063 | 14,263 |  | 10,348,117 | 14,271 |  | 9,938,198 | 12,523 |  |
|  | **#5** | 12,761,346 | 19,829 |  | 13,243,991 | 14,099 |  | 14,129,912 | 20,066 |  | 13,591,011 | 19,157 |  |
| **Water** |  | 3,789,807 | 0 | **N/A** | 3,864,601 | 0 | **N/A** | 4,517,577 | 0 | **N/A** | 4,583,998 | 0 | **N/A** |

*ID#: cell controls extracted, and sequenced at different time-points. EBV: Epstein-Barr virus, N/A Not applicable, ppm: reads per million of total classified reads*
