## Supplementary figures and images for "Optimization of cerebrospinal fluid microbial metagenomic sequencing diagnostics"

### Additional Figure 1

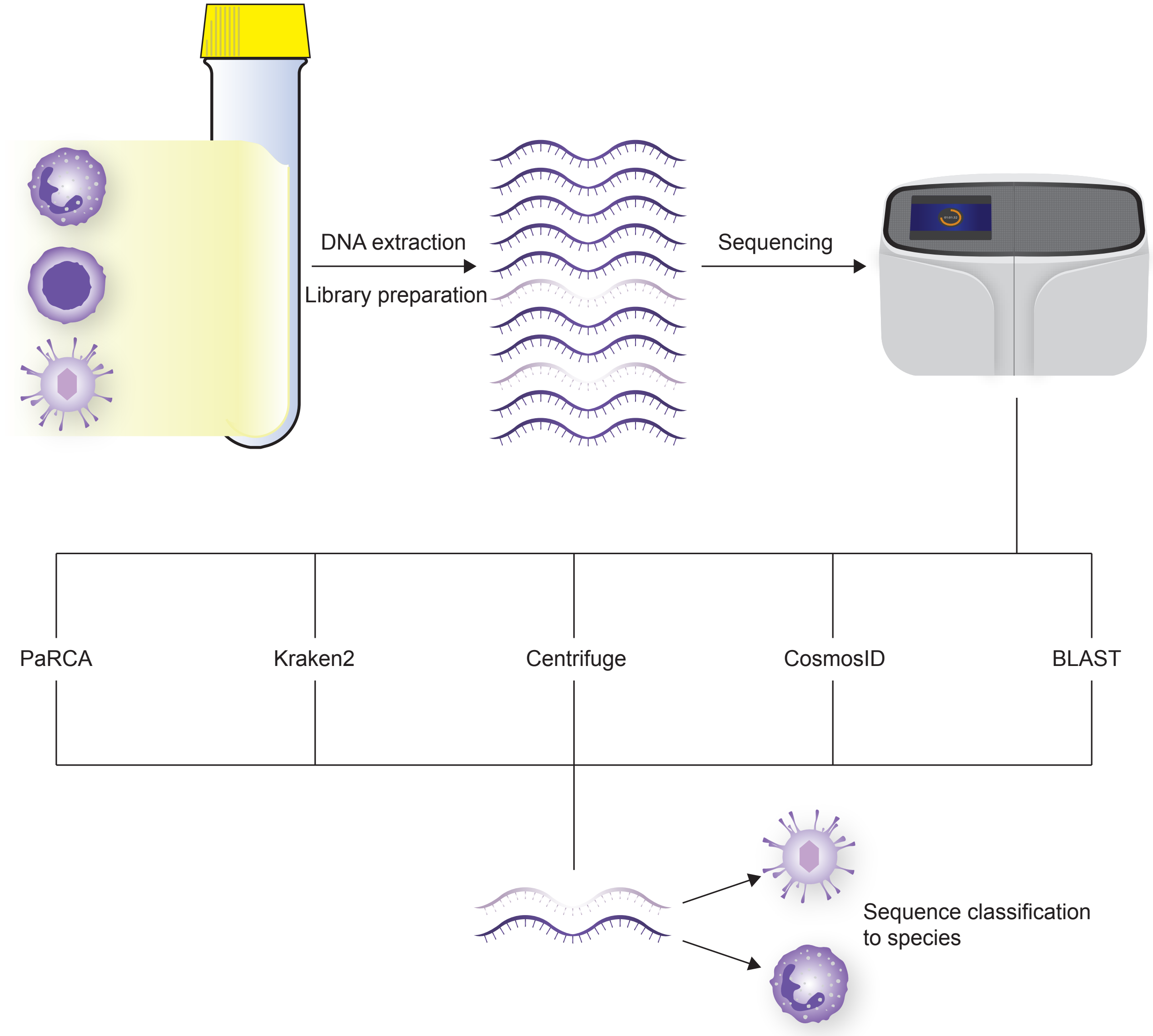

### Additional Figure 3

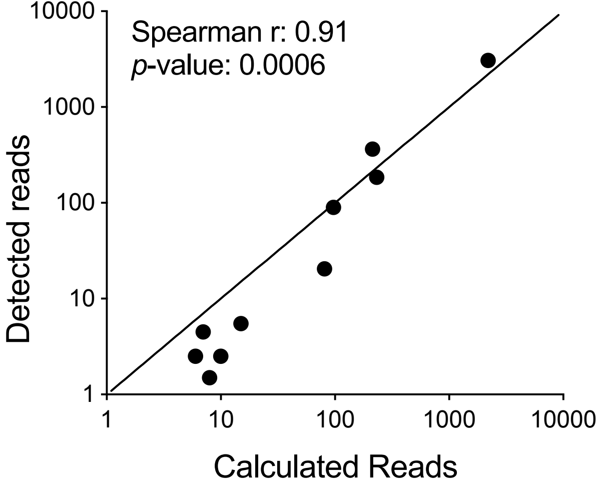

### Additional Figure 4

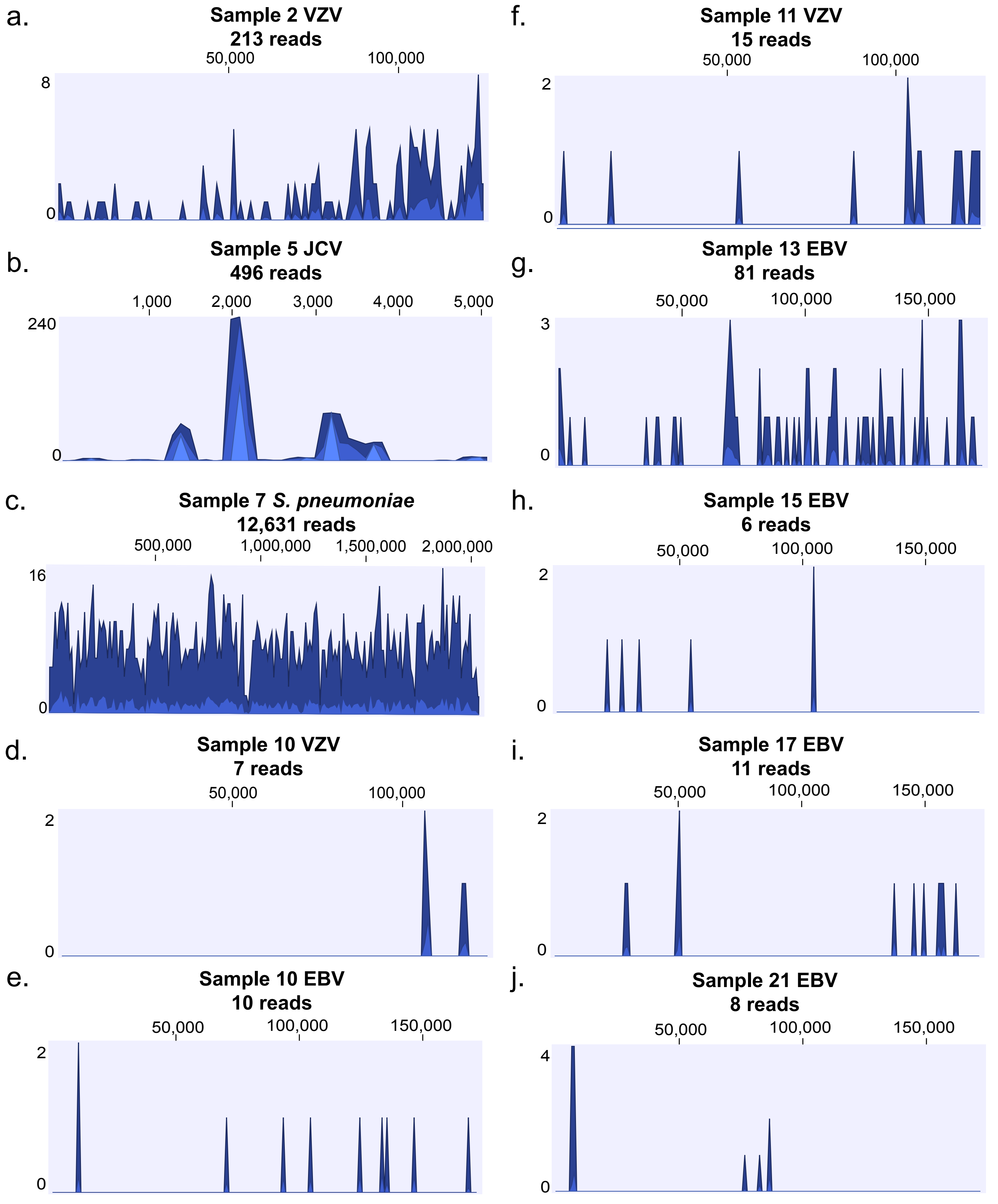

### Additional Figure 5

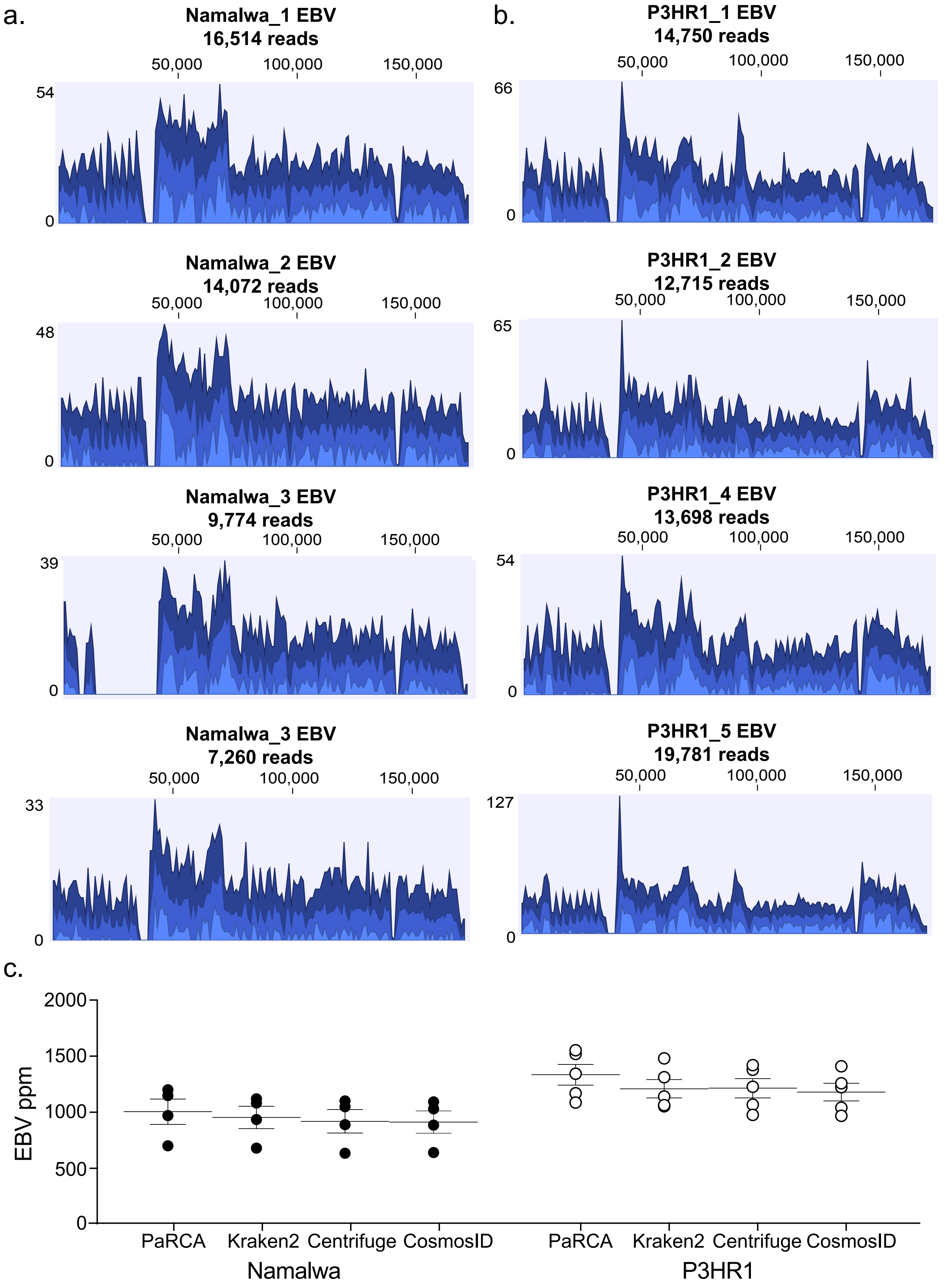

### Additional Figure 6

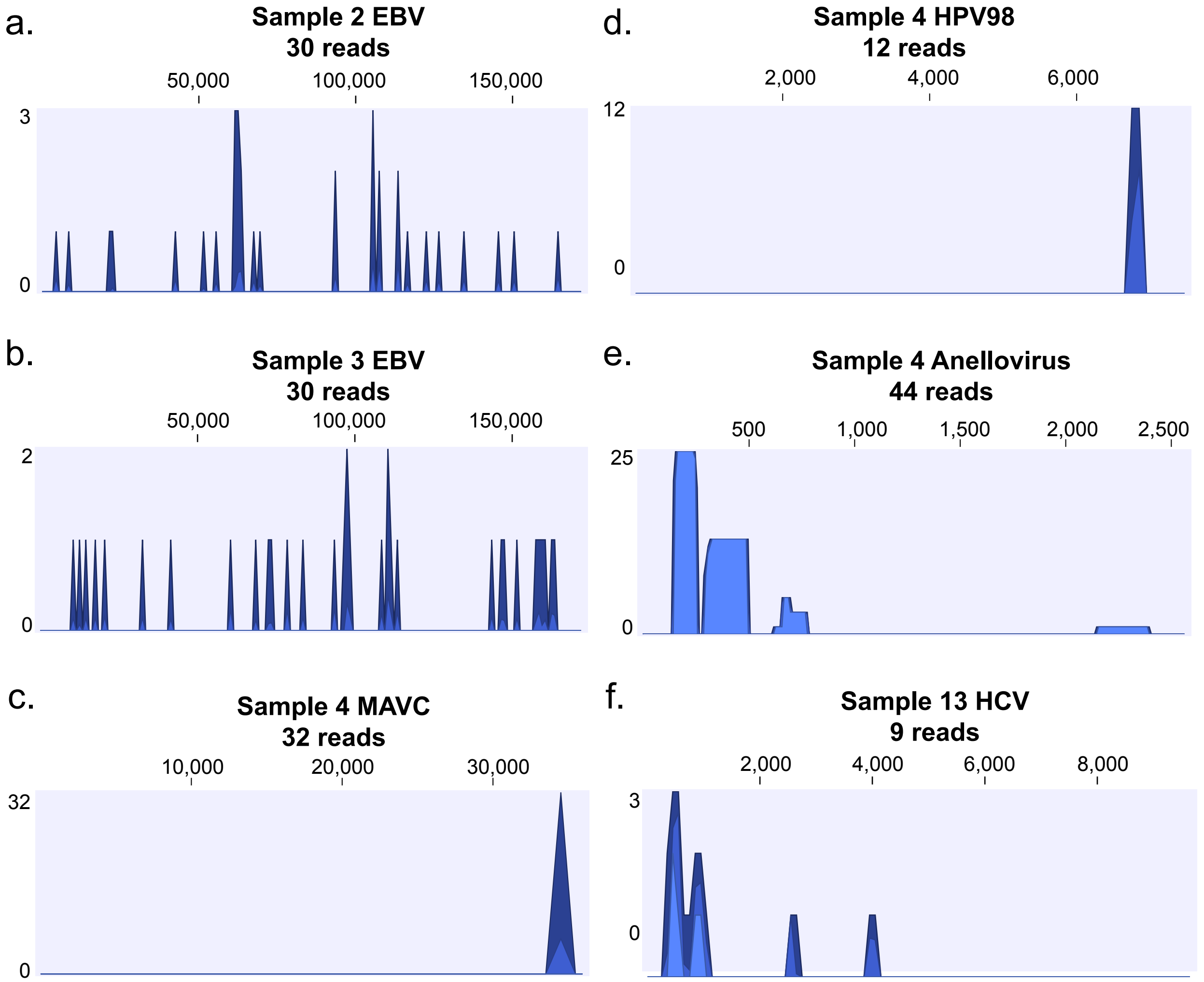
