## Additional Figure 2 for "Optimization of cerebrospinal fluid microbial metagenomic sequencing diagnostics"

### Sample 8 - 1,455,168 total classified reads

| Viruses | - | | |
|---|---|---|---|
| Hits | Taxid | % (RNA) | Reads (RNA) |
| | | >0.01 | |
| Escherichia virus T7 | 10760 | 0.0584 | 1101 |
| Enterobacteria phage T4 sensu lato | 348604 | 0.0135 | 254 |
| Enterovirus A | 138948 | 0.0119 | 224 |
| Enterovirus B | 138949 | 0.0316 | 595 |

### Sample 9 - 1,875,002 total classified reads

| Viruses | - | | |
|---|---|---|---|
| Hits | Taxid | % (RNA) | Reads (RNA) |
| | | >0.01 | |
| Enterobacteria phage T4 sensu lato | 348604 | 4.2333 | 61783 |
| Enterovirus B | 138949 | 0.1643 | 2398 |
| Escherichia virus M13 | 1977402 | 0.1433 | 2092 |
| Enterobacteria phage RB3 | 31533 | 0.0843 | 1231 |
| Escherichia virus T7 | 1985738 | 0.0276 | 403 |
| Pseudomonas phage PPpW-4 | 1279083 | 0.0246 | 359 |
| Shigella phage SHFML-26 | 1863009 | 0.0166 | 242 |
